# Molecular organization of integrin-based adhesion complexes in mouse Embryonic Stem Cells

**DOI:** 10.1101/416503

**Authors:** Shumin Xia, Evelyn K. F. Yim, Pakorn Kanchanawong

## Abstract

The mechanical microenvironment serves as an important factor influencing stem cell differentiation. Mechanobiological responses depend strongly on actomyosin contractility and integrin-based cell-extracellular matrix (ECM) interactions mediated by adhesive structures such as focal adhesions (FAs). While the roles of FAs in mechanobiology have been intensively studied in many mesenchymal and migratory cell types, recently it has been recognized that certain pluripotent stem cells (PSCs) exhibited significantly attenuated FA-mediated mechanobiological responses. FAs in such PSCs are sparsely distributed and much less prominent in comparison to “classical” FAs of typical adherent cells. Despite these differences, insights into how FAs in PSCs are structurally organized to perform their functions are still elusive. Using mouse embryonic stem cells (mESCs) to study PSC-ECM interactions, here we surveyed the molecular composition and nanostructural organization of FAs. We found that despite being small in size, mESC FAs appeared to be compositionally mature, containing markers such as vinculin, zyxin, and α-actinin, and dependent on myosin II contractility. Using super-resolution microscopy, we revealed that mESC FAs were organized into a conserved multilayer nanoscale architecture. However, the nanodomain organization was compressed in mESCs, with the force transduction layer spanning ∼ 10 nm, significantly more compact than in FAs of other cell types. Furthermore, we found that the position and orientation of vinculin, a key mechanotransduction protein, were modulated in an ECM-dependent manner. Our analysis also revealed that while most core FA genes were expressed, the expression of LIM domain proteins was comparatively lower in PSCs. Altogether our results suggest that while core structural and mechanosensitive elements are operational in mESC FAs, their structural organization and regulatory aspects may diverge significantly from “classical” FAs, which may account for the attenuated mechanobiological responses of these cell types.

## Introduction

Pluripotent stem cells (PSCs), such as embryonic stem cells (ESCs) and induced pluripotent stem cells (iPSCs), are capable of differentiation into virtually any cell types in the body and of unlimited self-renewal. Nevertheless, despite its numerous clinical potentials, the therapeutic applications of PSCs are, to a significant extent, held back by the incomplete understanding of factors governing their cell-fate specification and cell maturation^1^. Over the past few decades, mechanical cues have been found to be indispensable for many biological processes, including developmental morphogenesis, wound healing, and stem cell differentiation^2-3^. This has prompted intensive interest in the modulation of substrate mechanics or perturbation of cell contractility pathways for stem cell engineering^3^. Integrin-mediated adhesion to the extracellular matrices (ECM) together with the actomyosin contractility apparatus play essential roles in cellular sensing of the mechanical microenvironment^4^. However, much of our current insight into integrin-mediated mechanotransduction has been based on studies performed in specialized adherent cells (e.g. fibroblasts) or adult stem cells (e.g. mesenchymal stem cells)^5-7^, and whether these mechanisms are immediately translatable to PSCs remains to be validated.

The engagement of integrin mobilizes numerous structural and signaling proteins to form integrin-based adhesion complexes which serve as structural and mechanical links with the actin cytoskeleton. Specific terminologies for different maturation states of such structures include nascent adhesion (NA), focal complex (FC), focal adhesion (FA) and fibrillar adhesion (FB)^8^. These organelles are also collectively called FAs as henceforth referred to in this study. Mechanical force impinging upon each FA results in the elongation, enlargement, and compositional remodeling, a process termed FA maturation^9-10^. NAs arise first at the leading edge as diffraction-limited complexes. A small fraction of NAs then grow in size and mature into FCs and FAs in turn, concomitantly with the turnover of the remainder^11-12^. FAs transmit actomyosin-generated traction force against ECM and participate in a myriad of signaling pathways, and thus play integral roles in cell migration, ECM remodeling, and substrate-rigidity sensing^13-15^.

Recent studies indicated that FAs in mESCs and human ESCs (hESCs) are sparsely distributed and much less prominent compared to the well-characterized “classical” FAs of adherent specialized or mesenchymal cells^16-17^. Earlier studies^16-18^ have shown that PSCs require ECM adhesion to maintain proliferation and pluripotency, but the strength of the adhesions to fibronectin and the level of actin cytoskeletal organization were significantly lower^19^. Notably, the cytomechanical properties of PSCs and their responses toward mechanical stimuli were also found to be significantly attenuated vis-a-vis differentiated cells^20-21^. These well-documented phenotypic differences raise the possibility that FAs maybe organized or regulated differently in PSCs. Given the intensive interest in PSC-ECM interactions^22-23^, a detailed characterization of FAs in PSCs at the compositional and ultrastructural level should be highly informative.

FAs consist of a highly diverse set of molecular components—its proteome is usually referred to as the integrin “adhesome” ^24^. Recent efforts to catalogue the adhesome have primarily focused on proteomic analysis of mesenchymal cells, however^25^. Meta-analysis of these proteomic datasets have identified the “consensus”, “core”, and “meta” adhesome components, with 10, 60, and 2412 proteins, respectively^25-26^. At the molecular-scale level, much of our current knowledge on the structural organization of FAs has been revealed by super-resolution or electron microscopy studies carried out on differentiated cells^27-30^. Distinct protein layers were observed to organize along the vertical (Z) axis perpendicular to the plasma membrane. These strata were originally defined as the integrin signaling layer (ISL; constituents: integrin cytoplasmic domain, paxillin and FAK), the force transduction layer (FTL; constituents: talin and vinculin), and the actin regulatory layer (ARL; constituents: zyxin, VASP and α-actinin). The latter of which also structurally integrate into the termini of the actin stress fiber. Such vertically stratified architecture appears to organize around talin, which forms a direct integrin-talin-actin force-transmitting connection spanning through the FAs and defines the central FTL nanodomain. However, to what extent are such nanostructural frameworks conserved in FAs of PSCs are not known.

In this study, we sought to characterize the molecular organization and nanoscale architecture of FAs in PSCs. To preclude the contribution of E-cadherin-mediated cell-cell junction which crosstalks with cell-ECM adhesion and participate in the regulation of cell and colony mechanics, we chose murine ESCs (mESCs) as a model PSC system due to its ability to survive and maintain pluripotency as single cells. In contrast, hESCs undergo rapid apoptosis when isolated into individual cells unless ROCK inhibitor is applied to interfere with cytoskeletal contractility, and hESCs are considered to correspond to the later epiblast stage^31^. From transcriptome analysis, we found that although a majority of adhesome genes are expressed, comparatively lower expression level of LIM domain proteins appeared to be a key characteristic of PSCs. We also corroborated the significant morphological and spatial organization differences between mESC FAs and the “classical” mesenchymal FAs. Nevertheless, despite such differences our results indicated that mESC FAs are sensitive to myosin II-dependent contractility and contained core FA proteins, including markers of mature FAs, suggesting that they are compositionally mature. Using super-resolution microscopy to measure the vertical (Z-axis) position of major FA proteins, we found that while mESC FAs exhibited the conserved multi-layer ISL/FTL/ARL nanoscale architecture across different ECMs, the FTL nanodomain is more compact and the nanoscale distribution and orientation of vinculin differ from previous observations in “classical” FAs in an ECM-dependent manner. Our data identified potential structural and molecular basis that may account for the attenuated mechanobiological responses of these cell types, through the modulation of FA components and their nanoscale organization.

## Materials and Methods

### Cell culture

The E14.1 mESCs were received from Dr. Cheng Gee Koh (Nanyang Technological University, Singapore). To culture these cells, the culture dishes were pre-coated with 0.1% (wt/vol) gelatin (G1890, Sigma) for overnight at 4 ^°^C. The cell culture media consisted of high glucose DMEM (10566016, Invitrogen), 15% ES quality FBS (16141079, Invitrogen), 1% MEM nonessential amino acids (1114005, Invitrogen), 1% sodium pyruvate (11360070, Invitrogen), 1% penicillin/streptomycin (15140122, Invitrogen) and 55 μM 2-Mercaptoethanol (21985-023, Invitrogen), with additional 10^3^ unit of leukemia inhibitory factor (LIF, ESG1107, Millipore) supplemented freshly. The culture media was changed every day and cells were passaged every 3 days. Mouse Embryonic Fibroblasts (MEF) were received as a gift from the laboratory of Dr. Alexander Bershadsky (Mechanobiology Institute, Singapore). For the culture of MEFs, the media used consist of high glucose DMEM, 10% FBS (10082147, Invitrogen), 1% sodium pyruvate and 1% penicillin/streptomycin. Cells were maintained at 37 ^°^C with 5% CO_2_. Mycoplasma contamination test was performed monthly.

### Cell transfection and expression vectors

Lipofectamine 2000 (11668019, Invitrogen) was used to transfect the E14.1 mESCs according to the manufacturer’s instruction. Fluorescent protein fusion constructs for EGFP-Paxillin, EGFP-Talin (N-terminal FP fusion), mEmerald-Talin (C-terminal FP fusion), EGFP-vinculin (N-terminal FP fusion), tdEos-vinculin (C-terminal FP fusion), EGFP-zyxin, mEos2-VASP were described previously^28-29^. EGFP-FAK was a gift from the laboratory of Dr. Alexander Bershadsky. EGFP-vinculin T12 (N-terminal FP fusion) was a gift from the laboratory of Michael Sheetz (Mechanobiology Institute, Singapore). EGFP-VD1 (vinculin 1-258) and EGFP-vinculin T12 (C-terminal FP fusion) were generated by the protein cloning expression core in Mechanobiology Institute. mEmerald-Integrin-Linked-Kinase (ILK) was generated in the laboratory of Michael W. Davidson (The Florida State University) and available via Addgene depository (#54126).

### ECM coating and sample preparation

Clean No. 1 coverslip (Paul Marienfeld), glass-bottom dishes (Iwaki, Japan), and silicon wafers were used as imaging substrates. Three ECM proteins were used for substrate surface coating, including 10 μg/ml fibronectin (F1141, Sigma), 50 μg/ml laminin (23017015, Gibco) and 0.1% (wt/vol) gelatin. For ECM-specific experiments, DMEM and ES quality FBS were substituted with KnockOut™ DMEM (10829018, Gibco) and KnockOut™ Serum Replacement (10828010, Gibco) respectively to rule out the deposition of mixed ECM proteins from the serum. Cells were fixed after 6 h seeding for further staining. For pharmacological perturbations, cells were treated with 10 μM Y-27632 (Y0503, Sigma) or 10 μM Blebbistatin (B0560, Sigma) for 45 min before fixation.

### Cell fixation and immunofluorescence staining

Cells were fixed by pre-warmed 4% paraformaldehyde (PFA, Electron Microscopy Sciences) diluted in PHEM buffer (60 mM PIPES, 25 mM HEPES, 10 mM EGTA, and 2 mM MgSO_4_, pH 7.0) for 10-15 mins at room temperature. For permeabilization, 0.1% Triton X-100 in phosphate buffered saline (PBS) was applied to cells for 3 mins. For immunofluorescence staining, cells were first blocked by 4% bovine serum albumin (BSA, A7906, Sigma) in PBS for 1 h, followed by the incubation of primary antibodies in 1% BSA for overnight at 4 ^°^C. The samples were washed thrice with PBS before applying secondary antibodies in 1% BSA with or without dye-conjugated phalloidin at room temperature for 45 min.

Primary antibodies used for immunofluorescence were rat anti-CD29 (Integrin β1, 553715, BD), rabbit anti-Paxillin (SC-5574, Santa Cruz), mouse anti-Vinculin (V4505, Sigma), mouse anti-Talin (T3287, Sigma), mouse anti-Zyxin (ab58210, Abcam), mouse anti-FAK (610088, BD), mouse anti-p-Tyrosine (05-1050X, Millipore), mouse anti-α-actinin (A5044, Sigma), rabbit anti-myosin IIA (M8064, Sigma), mouse anti-Oct4 (MAB4419, Millipore) and rabbit anti-Nanog (sc33759, Santa Cruz). Secondary antibodies, phalloidins and DAPI were all from Invitrogen: Alexa Fluor 488 Goat anti-mouse IgG (A10680), Alexa Fluor 568 Goat anti-mouse IgG (A21124), Alexa Fluor 488 Goat anti-rabbit IgG (A11008) and Alexa Fluor 568 Goat anti-rabbit IgG (A11011). F-actin was stained by Alexa Fluor 488 Phalloidin (A12379) or Alexa Fluor 568 Phalloidin (A12380). The nuclei were stained by DAPI (D1306).

### Polyacrylamide gel (PAAG) substrate fabrication

Soft PAAG substrates were generated following a protocol described previously^32^. In brief, various concentration of Acrylamide (161-0140, Bio-rad) and Bisacrylamide (161-0142, Bio-rad) solution were mixed to achieve different stiffness as follows: 0.6 kPa (5% Acry/0.075% Bis), 2.3 kPa (7.5% Acry/0.075% Bis), 8.6 kPa (7.5% Acry/0.3% Bis), 30 kPa (12% Acry/0.28% Bis) and 55 kPa (12% Acry/0.6% Bis). To polymerize the PAAG, 10% Ammonium persulfate and TEMED were mixed with Bis-Acrylamide solution in de-ionized water. The solution was then immediately applied to a hydrophobic glass slide, and covered with an aminosilanized 25 mm diameter glass coverslip. After about 10 min polymerization, the gel with the 25 mm coverslip was carefully peeled off from the slide. The PAAG surface was then crosslinked with sulfo-SANPAH (22589, Thermo Scientific) under 365 nm UV activation, followed by fibronectin coating at concentration of 1mg/ml overnight at 4°C.

### Total Internal Reflection Fluorescence (TIRF) microscopy

TIRF imaging and surface-generated structured illumination microscopy were performed on a Nikon Eclipse Ti inverted microscope (Nikon Instruments, Japan), equipped with a motorized TIRF illuminator, an ORCA-flash 4.0 sCMOS camera (Hamamatsu), a light emitting diode (LED)-based epifluorescence excitation source (SOLA, Lumencor), and a CO_2_ and temperature control chamber (Okolab). In addition, a laser combiner (100 mW 405 nm, 60 mW 488nm, 50 mW 561 nm, and 100 mW 642nm solid-state lasers, Omicron Laserage) was coupled with a polarization-maintaining optical fiber. The objective used was a 60X N.A 1.49 Apo TIRF objective lens (Nikon Instruments).

### Spinning-disk confocal microscopy

For cells seeded on polyacrylamide gel, imaging was performed using Nikon Eclipse Ti inverted microscope equipped with a CSU-W1 spinning disk confocal unit (Yokogawa Electric Corporation), a ProEM-HS 1024B eXcelon3 EMCCD camera (Princeton Instrument), a laser combiner (100 mW 405 nm, 150 mW 488 nm, 100 mW 561 nm, and 100 mW 642 nm solid-state lasers) and a 100 X NA 1.45 oil objective lens (Nikon Instruments, Japan).

### Surface-generated structured illumination microscopy

Measurement of protein position along z-axis with nanometric resolution was performed using surface-generated structured illumination microscopy techniques (VIA-FLIC: Variable Incidence Angle Fluorescence Interference Contrast, or SAIM: Scanning Angle Interference Microscopy) described previously ^33-34^. Briefly, 4-inch silicon wafers (Bonda Technology) containing ∼500 nm thermal SiO2 layer were used as the imaging substrate. The thickness of the thermal oxide on each wafer was measured with sub-nanometer precision using a UV-visible variable angle spectroscopic ellipsometer (UV-VIS-VASE, J.A. Woollam) at A-STAR, IMRE, Singapore. Each wafer was cut into 1×1 cm^2^ pieces using a diamond-tip pen, followed by sequential cleaning using acetone and 1M KOH with sonication for 20 mins each. The wafers were then chemically activated by 0.5% (vol/vol) (3-aminopropyl) trimethoxy-silane (Sigma) in MilliQ water and 0.5% (vol/vol) glutaraldehyde (Electron Microscopy Sciences) in PBS for 1 h each, with 5 times x 5 minutes sonication in milliQ water after each step. Finally, the wafers were dried by nitrogen gas and sterilized by UV.

Proteins of interest were imaged using fluorescent protein fusion constructs transfected into the mESCs. Transfected cells were cultured on ECM-coated silicon wafer and fixed as described above. The samples were mounted with the cell side facing downward in a 27 mm glass bottom dish (# 3910-035, Iwaki) filled with PBS. A thumb screw was placed atop the wafer for neutral buoyancy. For each measurement, a series of raw images was acquired at 15 incident angles (from 0° to 56° with 4° increment). Focal adhesions region of interests (ROIs) were defined by thresholding or Otsu-based segmentation. The z-position for each ROI pixels were then computed using IDL-based custom-written software described previously^29, 35^. The median Z-position value of each ROI was used as the representative protein positions. Topographic images of protein z-position were generated using color to encode the z-position.

### Morphometric quantification of cells and FAs

Cell size and focal adhesion size was quantified on an IDL-based custom-written software (https://github.com/KanchanawongLab). Cell ROI masks were generated based on actin channel, while the FA masks were generated using paxillin channel. To generate the FA mask, the images were first background-subtracted using the Sternberg rolling-ball algorithm^36^ and the threshold for FA was set manually determined. Manual curation was also performed to exclude segmented non-FA regions.

### Transcriptome Analysis

RNAseq data for mouse iPSC, mESC, and MEF was retrieved from DBTMEE database^37^ (http://dbtmee.hgc.jp/index.php). For mESC (E14.1), hESC, and HT1080, RNAseq data was retrieved from Project ENCODE^38^ (https://www.encodeproject.org/) with the following dataset # ENCSR000CWC, ENCSR537BCG, ENCSR535VTR, respectively. For analysis, Log_2_ of the replicate-averaged of transcript quantifications in FPKM (Fragments per kilobase million) units were used. Clustering and correlation analysis were performed in R.

### Statistical Analysis

Statistical analysis and graph plotting were carried out using the following software: OriginPro, GraphPad Prism and R. Unpaired student’s t-test was performed for all the significance testing, unless otherwise specified. Details on data presentation are described in the legends of each figure.

## Results

### Expression Profiles of Core Integrin-based Adhesome in mESCs

The sparsity and small size of FAs in mESCs raised a question of whether mESC FAs may differ in composition from the “classical” FAs observed in adherent specialized cells. Recently, whole transcriptome RNA sequencing (RNAseq) has become widely accessible, with transcription profiles of a wide range of cell types available in public databases^39^. Although RNAseq data does not report on the enrichment of proteins or gene-products in specific sub-cellular compartments such as FAs, it nevertheless can be highly informative on the absence or abundance of given transcripts. Hence we used the core and meta-adhesome gene lists reported in earlier proteomic studies to investigate the expression of FA proteins in mESCs. For comparison between PSCs and fibroblastic cells, we retrieved the RNAseq data of mESCs, mouse iPSCs, and mouse embryonic fibroblast (MEF) from DBTMEE database^37^, as well as validated data on mESCs (E14.1), hESCs and fibrosarcoma cell line HT1080 released by Project ENCODE^38^. For the core adhesome list, we have chosen 56 out of 60 genes for analysis (Table S1) while excluding RPL23A (60S Ribosomal protein L23a), FAU (40S Ribosomal protein S30), PPIB (Peptidyl-prolyl Cis-Trans Isomerase B), and P4HB (Protein Disulfide Isomerase), which are ubiquitous housekeeping proteins expressed at very high levels in these cells. Hierarchical clustering of the expression level of the core adhesome genes is shown in Fig. 1. For the meta-adhesome, out of the total of 2412 genes, 1999 genes were chosen for analysis (Table S2). We excluded genes whose expression were not detected in at least one cell types, Table S3) and housekeeping genes with extremely high expression levels (primarily ribosomal proteins, Table S4). The correlation plot of the remaining 1999 genes is shown in Fig. S1.

**Figure 1.**
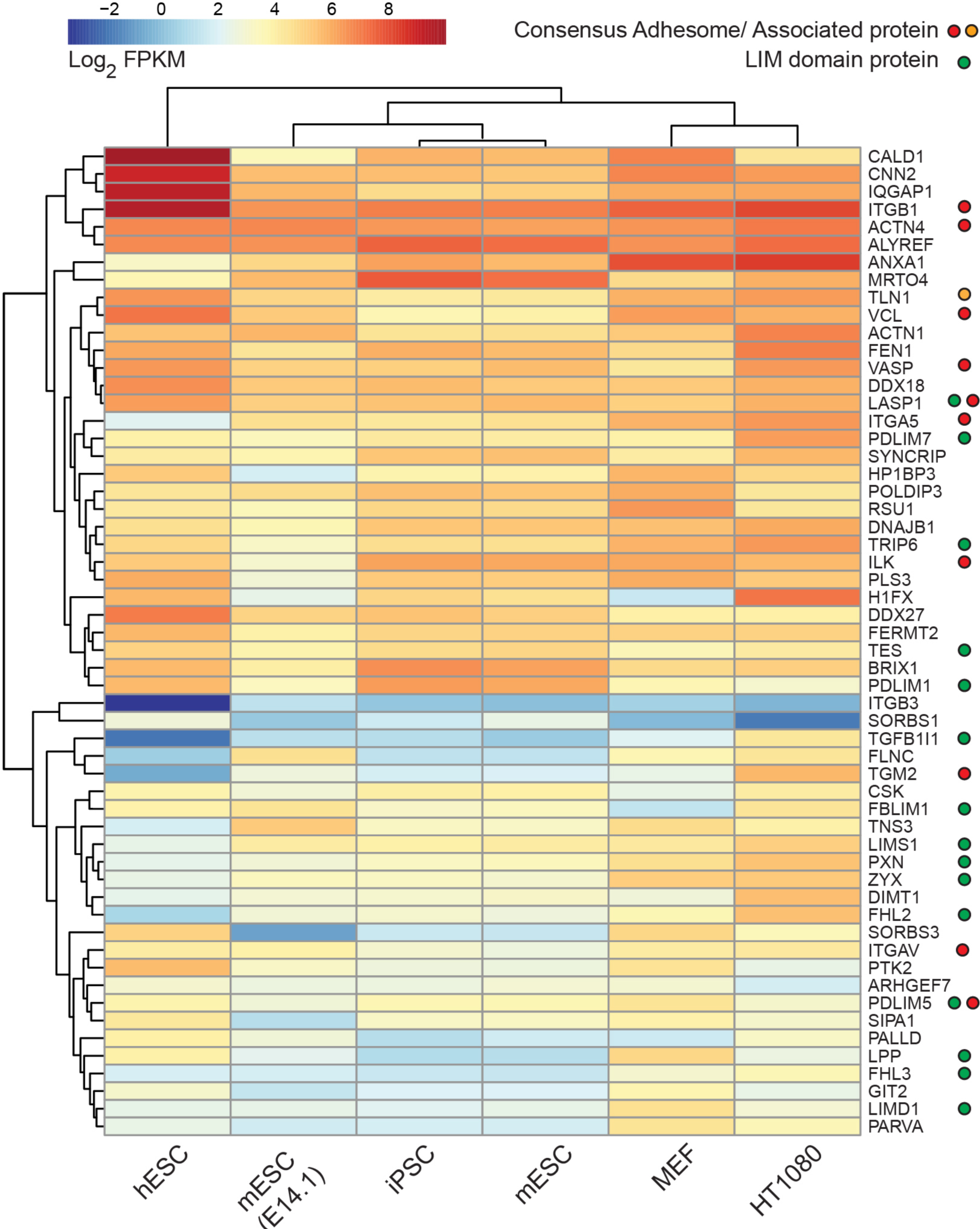
Expression Profiles of Core Integrin-based Adhesome in Pluripotent Stem Cells and Specialized Cells. Hierarchical clustering analysis of 56 gene expression profiles from RNA-seq data of four pluripotent stem cell lines (hESC, mESC(E14.1), iPSC, mESC) and two specialized cell lines (MEF, HT1080) were shown as heatmap and dendrogram. Data were retrieved from DBTMEE and ENCODE database. Log_2_ FPKM value was used for the analysis comparison. Cluster analysis was performed for both genes and cell lines using the complete linkage method based on Euclidean distance. Green circles indicate the LIM domain proteins, red circles indicate the consensus adhesome and yellow circle indicates the consensus adhesome-associated proteins.

As expected, a high degree of correlation in adhesome gene expression was found between the PSCs, which cluster together (Fig. 1). Of these, mESC and mouse iPSC from DBTMEE database of large-scale experiments exhibit highly similar expression profile. Notably, these also cluster with E14.1 (ENCODE) which corresponds to a different mESC line, but that which we used in this study as described further below^40^. For hESCs, the expression pattern retain significant similarity to mESCs and mouse iPSCs, albeit with a greater degree of difference as expected from their later epiblastic origin^41^. Likewise, the expression patterns of MEF and HT1080 cluster together and are distinct from the PSCs. Comparing “consensus” adhesome genes (red circle, Fig. 1) between the PSCs and the fibroblastic cells, we noted that PSCs exhibit somewhat lower expression levels (Log_2_ of FPKM) of α-actinin-4 (ACTN4), integrin β_1_ (ITGB1), integrin α_5_ (ITGA5), PDZ and LIM domain 5 (PDLIM5), transglutaminase-2 (TGM2), vinculin (VCL), and its close binding partner, talin-1 (TLN1), while relatively similar expression levels are observed for vasodilator stimulated phosphoprotein (VASP), integrin-linked kinase (ILK), integrin α_v_ (ITGAV), and LIM and SH3 domain protein1 (LASP1). Interestingly, our analysis also revealed that about one-third of the core adhesome genes (lower left corner, Fig. 1) were expressed at comparatively lower levels in PSCs. This cluster appears to be enriched in LIM domain proteins (green circles) such as paxillin (PXN), zyxin (ZYX), PDZ and LIM domain 5 (PDLIM5), LIM-Type Zinc Finger Domains 1 (LIMS1) or PINCH, Four-and-a-half LIM Domain 2 (FHL2), Four-and-a-half LIM Domain 3 (FHL3), LIM domain containing protein 1 (LIMD1), and Filamin Binding LIM Protein 1 (FBLIM1), Lipoma-Preferred Partner (LPP), and Transforming Growth Factor Beta-1-induced Transcript 1 (TGFB1I1). Altogether our analysis suggested that while FA components are generally expressed at substantial levels (Table S2), a notable reduction in the expression of LIM domain proteins appear to be a common signature for PSCs and may point to an interesting direction for further studies on PSC mechanotransduction.

### Responses of mESC FAs to substrate rigidity

Indeed, previous proteomic studies on “classical” FAs have identified LIM domain proteins as a significant portion of the contractility-dependent adhesome (proteins which were depleted from FAs upon contractility inhibition)^42^. This raised the possibility that the lowered expression of LIM domain proteins observed herein (Fig. 1) may affect the mechanosensitivity of mESC FAs. We therefore sought to address this by investigating the spatial organization of FAs in mESCs. Previously reported studies on ESC FAs, however, have largely focused on ESC colony^18, 43^. However, cell-ECM interactions in PSC colony are intrinsically heterogenous due to factors such as relative positions of cells within the colony (e.g., edge vs. center), colony geometry, while E-cadherin-mediated cell-cell junctions are also key contributors to cell mechanics. To investigate how ESCs interact with the ECM in the single cell state, we therefore dissociated the mESCs (E14.1^40^) and seeded them on fibronectin substrate as single cells and imaged at 6 h post-plating. To ascertain the pluripotency of these mESCs under such experimental condition, we probed the pluripotency marker Oct4 and Nanog by immunofluorescence microscopy^44^. The Oct4 and Nanog signals were observed primarily in the cell nucleus, as indicated by a co-staining by DAPI, suggesting that the cells retained their pluripotency (Fig. 2A).

**Figure 2.**
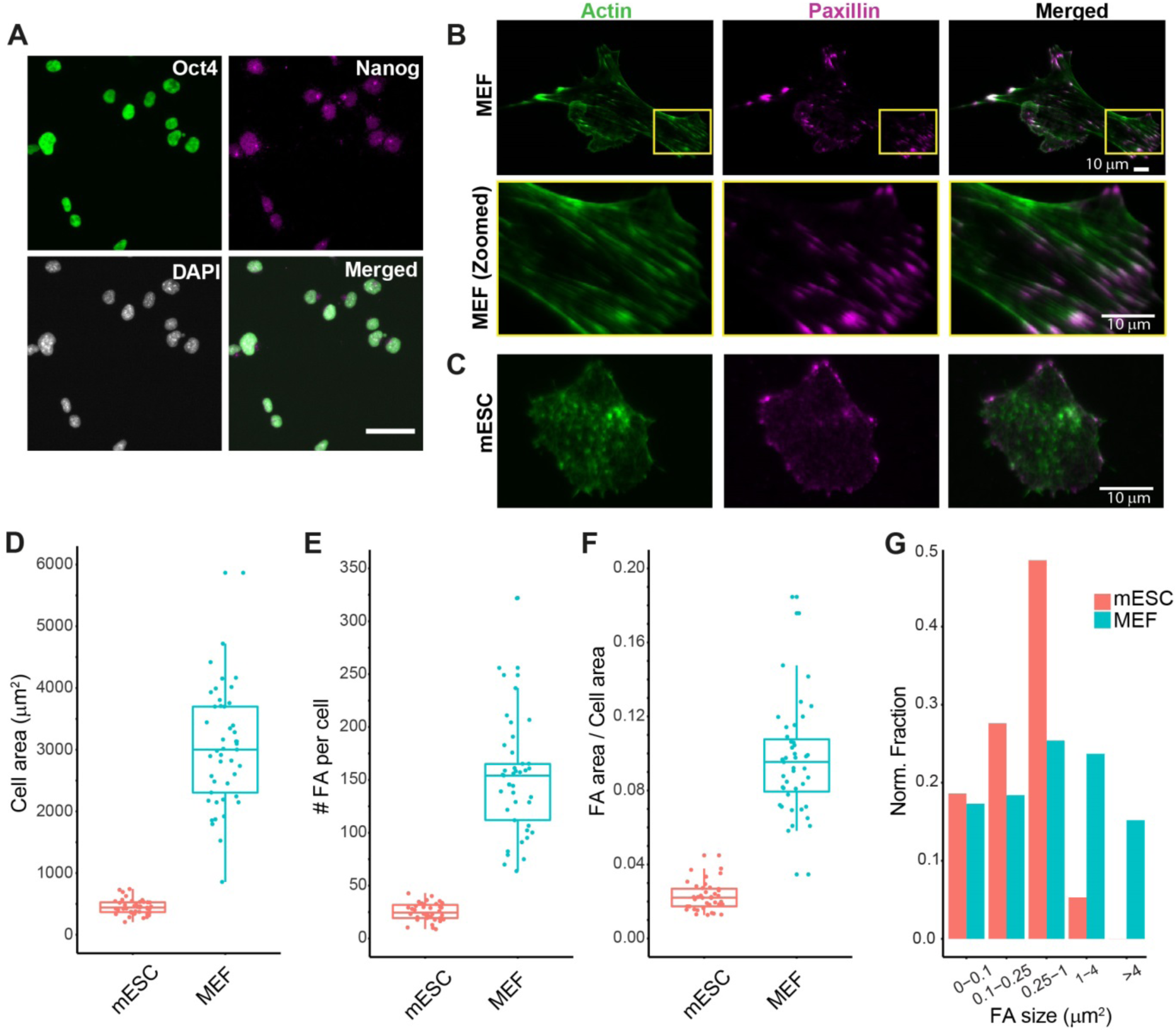
Comparison of Cell and FA morphometrics between mESCs and MEFs. **(A)** Immunofluorescence microscopy of the pluripotency marker Oct4 (green) and Nanog (magenta), together with the nuclei marker DAPI in mESCs after 6 h plating on FN. **(B, C)** Immunofluorescence microscopy of actin (green) and paxillin (magenta). Boxed region in the top row of **(B)** was magnified in the bottom row. Scale bar: 50 μm **(A)**, 10 μm **(B, C)**. **(D-G)** Quantifications on cell spreading area, FA area relative to the cell area, FA number in mESCs (blue) and MEF (red) were plotted using the box plot, with the middle line indicating the median value, the box representing the interquartile range, and the whiskers representing the 1.5X interquartile range. Each data point (n_cell-MEF_=43, n_cell-mESC_=40) was also co-plotted with the box. The FA size (n_FA-MEF_=6651, n_FA-mESC_=1052) was presented as a histogram with unequal bins.

A hallmark mechanobiological response of many adherent cells including fibroblasts and MSCs is the increase in cell spreading areas and FA sizes as a function of substrate stiffness^45-47^. To probe this effect in mESCs, we quantified the cell spreading area, FA size (using immunostained paxillin as FA marker), FA/cell area ratio, and FA number per cell in E14.1 mESCs, in comparison with mouse embryonic fibroblasts (MEFs), a representative specialized adherent cell (Fig. 2B-C). These cells were cultured on fibronectin-coated cover-glass or polyacrylamide gel (PAAG) with Young’s modulus of 0.6, 2.3, 8.6, 30, or 55 kPa, respectively. Our results showed that the average cell spreading area on fibronectin-coated glass of mESCs was around 450 μm^2^ compared to 3000 μm^2^ for MEF (Fig. 2D), while the FA size of mESCs was also relatively smaller than MEFs (Fig. 2G). Because of the much smaller cell size of mESCs, its FA density (both in terms of number and area) appeared to be much lower than MEF (Fig. 2E-F). In addition to the low density, FAs in mESCs were also primarily restricted to the cell periphery, as indicated by paxillin staining (Fig. 2C).

As shown in Fig. 3, we observed that in MEFs the cell spreading area increases in proportion to substrate rigidity, associated with the increasing prominence of FAs (Fig. 3B), demonstrating a robust mechanobiological response^48^. In contrast, for mESCs FAs remained small and confined to the cell periphery (Fig. 3A, images were shown as inverted contrast to aid in observing FAs; dark regions at cell centers correspond to the non-specific paxillin immunostaining background in the cell body). Furthermore, the cell spreading area of mESCs remained largely unchanged over the 0.6-55 kPa range, in stark contrast to MEFs (Fig. 3C, and Fig. S2). Altogether, our data corroborated earlier findings that the response of mESCs to substrate stiffness is significantly attenuated in comparison to mesenchymal cells^21^.

**Figure 3.**
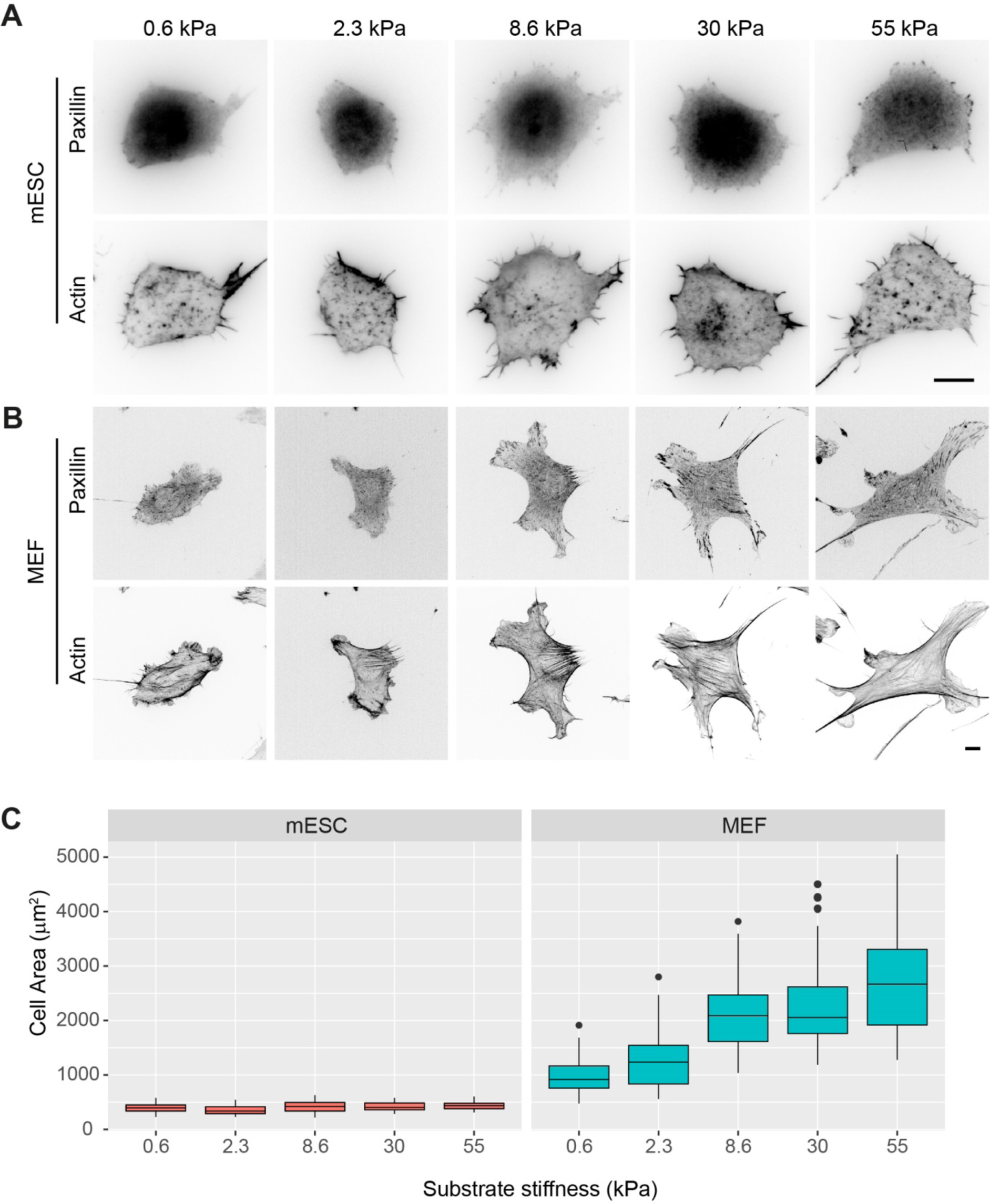
Responses of mESCs and MEFs to Substrate Rigidity. **(A, B)** mESCs (A) and MEF (B) plated on 0.6, 2.3, 8.6, 30, 55 kPa fibronectin-coated polyacrylamide gel (PAAG) were fixed and immunostained for paxillin and labeled by phalloidin. Scale bar: 10 μm. **(B)** Quantifications on cell spreading area from **(A)**. Data were presented as boxplot (n_cell-E14_=31, 34, 34, 32, 36 and n_cell-MEF_=33, 37, 37, 39, 34 for 0.6, 2.3, 8.6, 30, 55 kPa), with the whisker representing the range of 1.5x interquartile.

### FAs in mESCs are dependent on myosin II-contractility

The small size of mESC FAs raised further questions on whether these FAs are mature or mechanosensitive. Since FA maturation is commonly associated with myosin II-generated tension^10, 49^, we next examined the localization of endogenous myosin IIA in mESCs plated on different ECMs. We note that much of the existing literature on “classical” FAs involve fibronectin-dependent adhesions, whereas in studies on PSCs collagen-derived gelatin and laminin (basement membrane ECM) are commonly used. To study mESC responses to these ECMs, the substrates were coated with fibronectin, laminin, or gelatin, while serum-free media was used to preclude the involvement of serum-derived ECM. Cells were fixed 6 h post-plating and stained for myosin IIA, the pluripotency marker Oct4, and F-actin. As observed earlier (Fig. 2A), our results showed that the pluripotency marker was still retained and that the mESCs appeared to spread in a similar manner on fibronectin, laminin and gelatin (Fig. 4A). Myosin IIA was found to be largely restricted to cell periphery, likely associated with peripheral actin bundles, while the cell spreading areas (∼ 450 μm^2^) were not significantly different between different ECMs (Fig. 4B).

**Figure 4.**
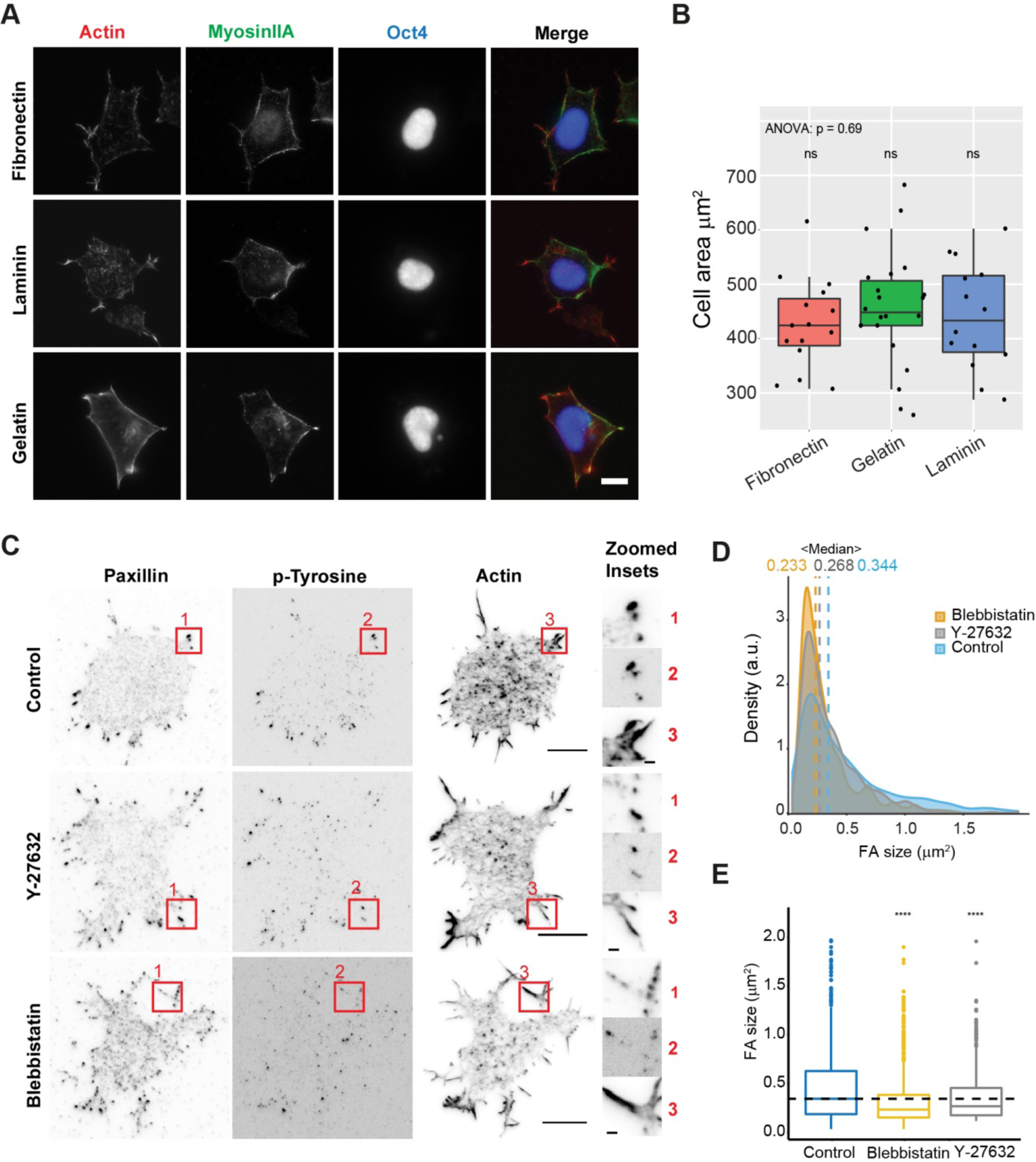
FAs in mESCs are dependent on myosin II-contractility (A) Immunofluorescence microscopy probing for Myosin IIA and Oct4, together with actin and **(B)** the quantification of the cell spreading area in mESCs plated on fibronectin, laminin and gelatin (n_cell-fibronectin_=15, n_cell-laminin_=14, n_cell-gelatin_=21). Data were presented as boxplot and data points, with the whisker representing the range of 1.5x interquartile. One-way ANOVA test was performed for the cell spreading area on three ECMs, and Student’s t-test between each condition to the average of the three conditions was performed for the significance level. ns: not significant. **(C)** Immunofluorescence microscopy probing for paxillin, phospho-tyrosine and actin on mESCs plated on the fibronectin-coated cover glass, untreated (control), treated with 10 μM Y-27632 and 10 μM blebbistatin (40 min), respectively. The boxed region was magnified in the right column. Scale bar: 10 μm (**A**, **C**); inset: 1μm (**C**). **(D, E)** FA size in control (blue), Y-27632 treated (grey) and blebbistatin treated (yellow) mESCs presented as density plot **(D)** and boxplot **(E)**. The median FA size was labeled as vertical dashed line, and the value is indicated on top of the density plot (n_FA-control_=572, n_FA-blebbistatin_=1524, n_FA-Y27632_=620). The dashed line in **(E)** represents the median FA size of control cells. Statistical hypothesis test was performed using Student’s t-test. ****: P<0.0001.

We next disrupted myosin II-generated contractility by applying either myosin II activity inhibitors, blebbistatin (10 μM, 40 min), or Rho kinase inhibitor, Y-27632 (10 μM, 40 min). Following immunofluorescence probing for paxillin (FA marker) and phosphor-tyrosine (FA signaling marker) and quantitative image analysis, we observed a significant reduction in both the sizes and FA signaling in comparison to control (Fig. 4C-E). Taken together, these results suggest that despite their relatively small morphology, low density, and sparse peripheral distributions, mESC FAs appear to be mechanosensitive and disassemble in response to the reduction in intracellular tension in a manner similar to “classical” FAs.

### Compositional maturation of mESC FAs on various ECMs

As the maturation of “classical” FAs involve the recruitment of key proteins such as vinculin and zyxin, we next sought to probe the composition of mESC FAs formed on different ECMs described above as these are recognized by different integrin heterodimers (fibronectin: α_5_β_1_ and α_v_β_3_, gelatin: α_1_β_1_ and α_2_β_1_, laminin: α_3_β_1_ and α_6_β_1_,). Cells were fixed 6 h post-plating and endogenous FA proteins were probed by immunostaining, with paxillin used as FA marker. As shown in Fig. 5A-C, we found that canonical proteins of mature FAs such as FAK, paxillin, talin, vinculin, zyxin, and α-actinin were distinctly localized to mESC FAs. In particular, the presence of zyxin and α-actinin, which are considered hallmarks of mature FAs in differentiated cells^50^, suggests that these three ECMs were all capable of supporting the compositional maturation of mESC FAs. Additionally, using monoclonal antibody against activated integrin β1 (clone 9EG7, prominent FA staining was observed for all three ECM conditions, indicative of integrin activation as expected^18^. The mESC FAs were also observed with strong phospho-tyrosine staining, signifying active signaling activities. Altogether, our results so far indicated that despite the attenuated morphological features and relatively subdued mechanosensitive responses, the mESC FAs are compositionally mature and sensitive to myosin II contractility.

**Figure 5.**
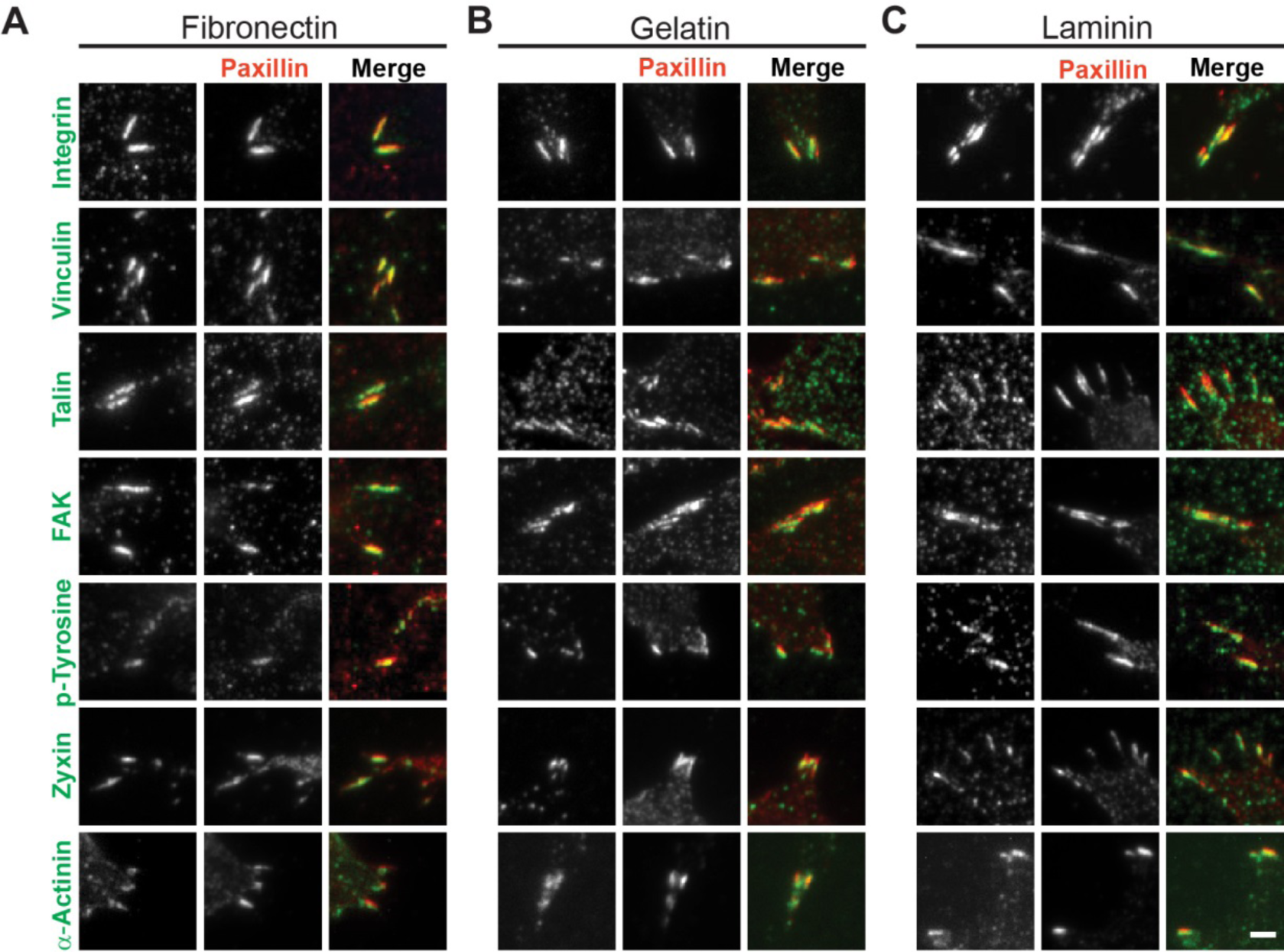
Compositional maturation of mESC FAs on various ECMs. **(A-C)** Zoom-in view of FAs in mESCs from immunofluorescence microscopy probing for endogenous integrin, vinculin, talin, FAK, phosphor-tyrosine, zyxin and α-actinin (green) together with paxillin (red) as a common marker in mESCs plated on fibronectin **(A)**, gelatin **(B)** and laminin **(C)**. Scale bar: 2 μm.

### Probing the Nanoscale Architecture of mESC FAs

Since FA force transmission is integral to integrin-mediated mechanosensitivity^4^, we next sought to map the molecular-scale organization of force transmission proteins and key adhesome components within mESC FAs. We used a surface-generated structured illumination techniques (SAIM: Scanning Angle Interference Microscopy) which enables ∼10-nm precision measurements of vertical (z)-position of fluorophores (Fig. 6A)^33-34^. As imaging probes, we used fluorescent protein (FP) fusion of FA proteins due to its <5-nm probe size suitable for the high precision of SAIM. As the nanoscale architecture of “classical” FAs have primarily been studied on fibronectin, we first investigated mESC FAs on fibronectin-coated substrate. For the measurement, mESCs expressing FP fusion probes were cultured for 6 h before fixation for imaging. The representative maps of the z-position of FA proteins relative to the surface of the imaging substrate (z=0) are shown in Fig. 6B. The median z-position (Z_median_) for each FA region-of-interest (ROI) is used as the representative z-positions for a given FA protein. The histograms and notched box plot for the distribution of Z_median_ are shown in Fig. 6C, along with the corresponding number of cells (blue) and number of FA ROIs (black), and the representative Z_median_ (red).

**Figure 6.**
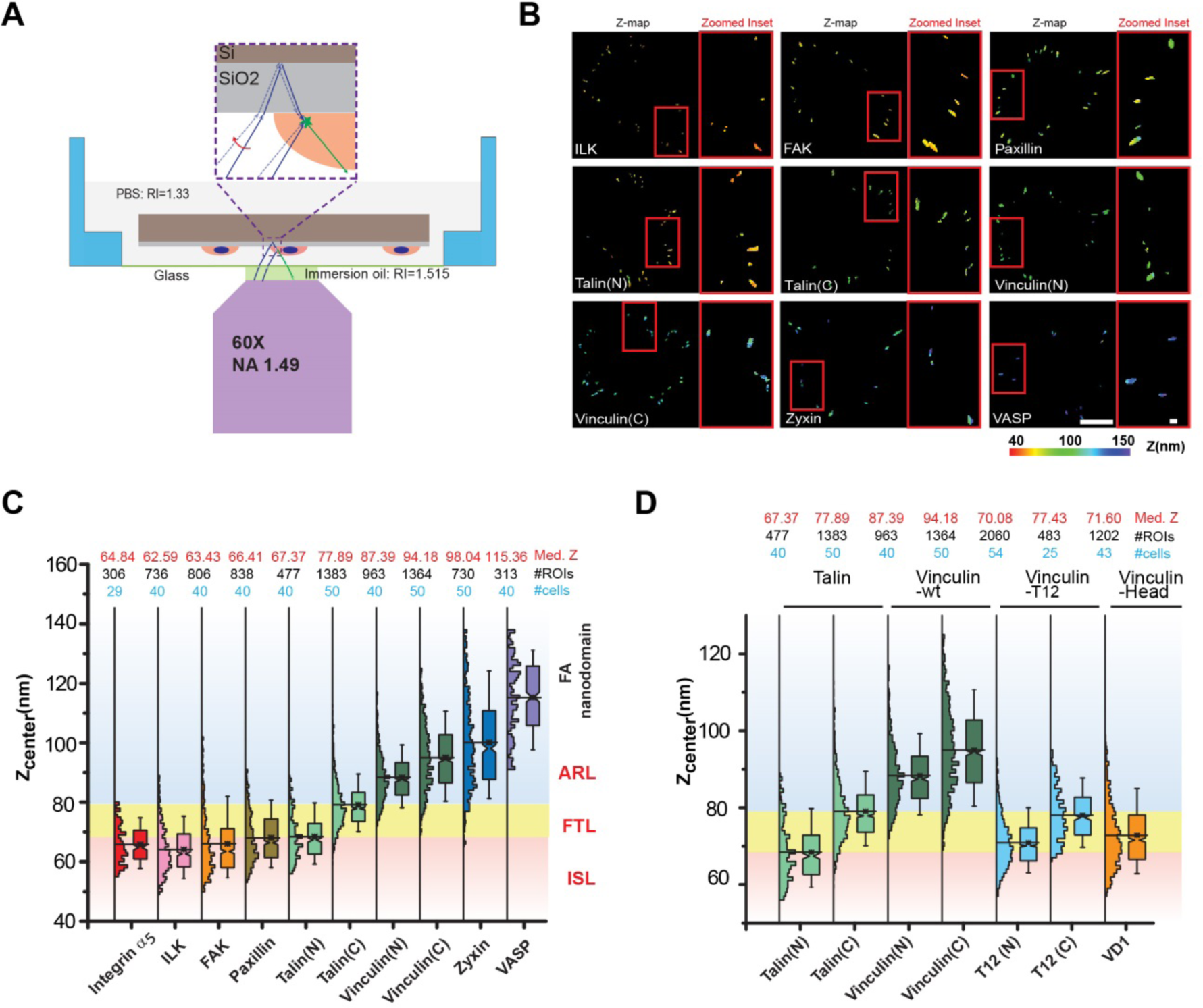
Probing the Nanoscale Organization of Proteins in mESC Fas. **(A)** Schematic of sample mounting and nanoscale Z-measurement using surface-generated structured illumination microscopy. **(B)** Topographic map of FA with color-encoded nanoscale Z position as measured by surface-generated structured illumination microscopy. The magnified inset region was shown to the right of each panel. Color bar indicates Z position relative to the surface. Scale bar: 10 μm; inset: 1 μm. **(C, D)** Nanoscale organization of canonical integrin-based adhesion proteins **(C)** and vinculin mutants **(D)** plotted as the histogram and notched box with whisker side by side using the Z-center of each FA ROI. The Z-position is relative to the surface. Numbers at the top of the plot indicate the median Z center (red), the number of ROIs (black) and the number of cells (blue). The red, yellow and blue regions indicate the integrin signaling layer (ISL), force transmission layer (FTL) and actin regulatory layer (ARL).

We determined the z-position of integrin α_5_ (FP label at C-terminal cytoplasmic domain) subunit of the fibronectin-binding integrin α_5_β_1_ to be at z = 64.84 nm. This also serves to outline the plasma membrane z-position. Together with ILK and FAK, paxillin was also observed in the vicinity of the integrin cytoplasmic domain, thus defining the ISL stratum as described previously ^51^. We next probed the organization of talin using fusion constructs with FPs fused at either the N- or C-terminus (Talin-N and Talin-C, respectively, Fig. 6C). We observed that the N-terminus of talin was at z = 67.37 nm, consistent with the N-terminal FERM domain engaging and activating integrin. In contrast, the C-terminal of talin was observed at a higher z-position, z =77.89 nm. Thus, talin in mESC FAs appears to be oriented in a uni-directional manner similar to in “classical” FAs of differentiated cells, although the span between the N- and C-terminus is much reduced, at about 10 nm, compared to in differentiated cells (25-30 nm) ^29, 51^. We also probed actin regulatory proteins such as zyxin, and VASP. These were observed at elevated z-positions (Zyxin, z = 98.04 nm; VASP, z = 115.36 nm), as expected from their close association with F-actin. From our measurements, we conclude that while FA proteins in mESC are stratified into distinct ISL/FTL/ARL nanodomains similar to previous observation in “classical” FAs, the boundary of these layers may need to be re-defined based on the measured protein positions. As shown in Fig. 6 and 7, the FTL is therefore defined as the interval between median z-position of N- and the C-terminal probes of talin (Talin-N and Talin-C), respectively.

**Figure 7.**
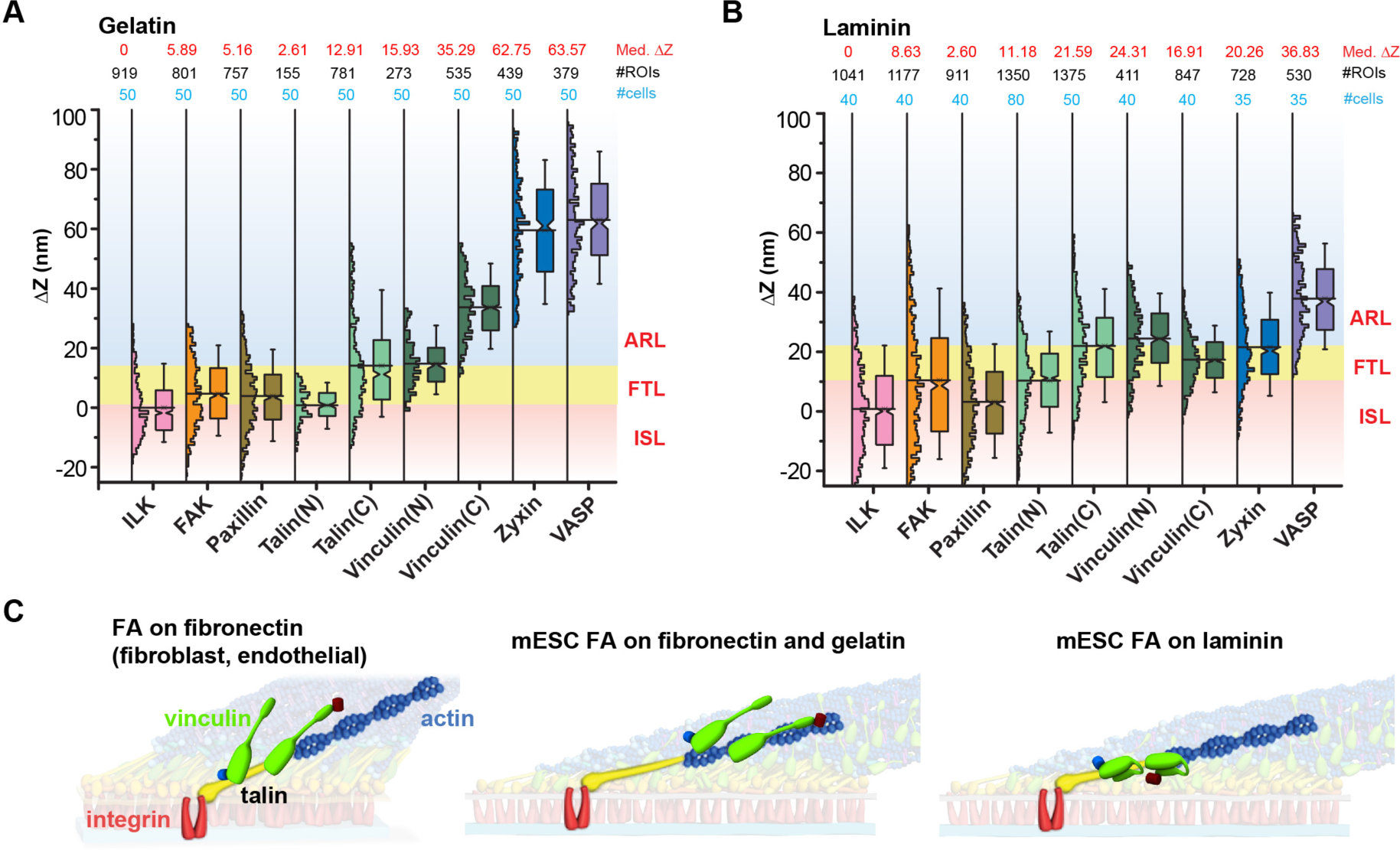
Nanoscale Architecture of mESC Fas. **(A, B)** Nanoscale organization of canonical integrin-based FA proteins in mESCs plated on gelatin **(A)** and laminin **(B)**. Data were plotted as histogram and notched box and whisker plot using Z-center of each FA ROI. Z-position is relative to the ILK as described in the main text. Numbers on top of the plot indicate the median Z center (red), the number of ROIs (black) and the number of cells (blue). The red, yellow and blue regions indicate the integrin signaling layer (ISL), force transmission layer (FTL) and actin regulatory layer (ARL). (**C**) Schematic diagram of the FA nanoscale architecture in mESCs, highlighting the core integrin-talin-actin connection (in red, yellow, blue, respectively). Vinculin is shown in green and marked by fluorescent protein probes at either the N- (blue) or C- (red) termini. Different configuration of vinculin in relation to integrin-talin-actin core is highlighted.

Next, we mapped the z-position of vinculin, using fusion constructs with FP fusion at the N- or C-terminus (vinculin-N and vinculin-C, respectively). Recent super-resolution studies on fibroblasts showed that while vinculin nanoscale distribution is centered in FTL, this can be re-distributed between ISL, FTL, and ARL through interactions with talin, F-actin, and phosphorylated paxillin, consistent with the notion of vinculin as the regulatable molecular clutch^52^. Surprisingly, our SAIM measurements revealed that in mESC FAs vinculin occupied relatively high z-positions, largely within ARL zone (vinculin-N: 87.39 nm; vinculin-C: 94.18 nm). Indeed, as the talin binding site of vinculin is located near the N-terminus, the minimal FTL distribution of vinculin-N (Fig. 6D, Fig. S3) suggest that a majority of vinculin may not physically localize with talin in mESC FAs on fibronectin substrate.

Vinculin is a conformationally autoinhibited multi-domain protein capable of force transmission in FAs via crosslinking between talin and F-actin. Strong autoinhibitory binding between the globular head domain (V_H_: residue 1-880) and the C-terminal domain (V_T_: residue 881-1067) can be relieved by binding to talin, or by site-specific mutations or domain truncation^53^. To further investigate the molecular basis of vinculin nanoscale distribution, we used an FP fusion of the VD1 talin-binding vinculin-head subdomain (residue 1-258) to map vinculin binding sites in mESC FAs. We found that the VD1 probe distinctly localize to the FTL at z = 71.6 nm, likely due to its binding to the middle span of talin rod domain (Fig. 6D, Fig. S3). We further corroborated this by making use of Vinculin-T12, a constitutively-active mutant of full-length vinculin^54^, with FP fusion at either the N- or the C-termini. We observed that the N-terminus of vinculin-T12 is located at z = 70.08 nm similar to VD1, while the C-terminus is at 77.43 nm (Fig. 6D). Altogether these results suggest that activated vinculin in mESC FAs can recapitulate the FTL nanoscale distribution of vinculin as observed in “classical” FAs, while also implying that wild-type vinculin in mESC FAs could be largely decoupled from talin, but may interact with ARL-localized protein instead. A potential ARL partner for vinculin is α-actinin which contain mechanosensitive cryptic vinculin binding site^55^. Further in-depth mechanistic dissection would be required for a better understanding of this process.

### Nanoscale Architecture of mESC FAs on Gelatin and Laminin

Since distinct integrin αβ heterodimers mediated adhesion to different ECMs, we also investigated whether the multi-layer architecture of mESC FAs are common across different ECMs. We carried out SAIM nanoscale mapping of FA proteins as above in mESCs cultured on laminin- or gelatin-coated Silicon wafer for 6 h. We observed that the absolute z-positions of FA proteins relative to the substrates are collectively shifted relative to fibronectin measurements, indicative of the differences in ECM thickness (Fig. S4). For example, ILK is locating at z= 113.7 nm on laminin and z= 69.06 nm on gelatin. To allow comparison across different ECMs, we therefore used the position of ILK as the reference position (Δz=0) (Fig. 7A-B, Table 1). Note also that the higher ECM thickness appeared to result in a somewhat reduced measurement precision as reflected by the broader distribution of the histograms (Fig. 7B), likely due to the reduced coherence of surface-generated structured illumination excitation field^56^.

**Table 1.** Organization of integrin adhesome proteins in mESCs as measured by surface-generated structured illumination microscopy. Z-position used are calculated relative to the median position of ILK to account for varying characteristic thickness of different extracellular matrices coatings as described in the main text.

| Protein | N<sub>FA</sub> | N<sub>cells</sub> | MedianZ<sub>FA</sub>(nm) | Mean Z<sub>FA</sub>(nm) | Std.dev.(nm) |
| --- | --- | --- | --- | --- | --- |
| ILK | 736 | 40 | 0 | 1.55 | 8.22 |
| FAK | 806 | 40 | 0.84 | 3.48 | 10.79 |
| Paxillin | 838 | 40 | 3.82 | 5.52 | 8.68 |
| Talin-N | 477 | 40 | 4.78 | 5.84 | 7.55 |
| Talin-C | 1383 | 50 | 15.30 | 16.52 | 8.03 |
| Vinculin-N | 963 | 40 | 24.80 | 25.75 | 8.43 |
| Vinculin-C | 1364 | 50 | 31.58 | 32.39 | 11.47 |
| Zyxin | 730 | 50 | 35.45 | 37.52 | 15.79 |
| VASP | 313 | 40 | 52.76 | 52.63 | 12.61 |

2) Laminin
| Protein | N<sub>FA</sub> | N<sub>cells</sub> | MedianZ<sub>FA</sub>(nm) | MeanZ<sub>FA</sub>(nm) | Std.dev. (nm) |
| --- | --- | --- | --- | --- | --- |
| ILK | 1041 | 40 | 0 | 0.83 | 15.37 |
| FAK | 1177 | 40 | 8.63 | 10.44 | 21.19 |
| Paxillin | 911 | 40 | 2.60 | 3.22 | 14.19 |
| Talin-N | 1350 | 80 | 11.18 | 10.32 | 13.04 |
| Talin-C | 1375 | 50 | 21.59 | 21.97 | 14.21 |
| Vinculin-N | 411 | 40 | 24.31 | 24.46 | 11.45 |
| Vinculin-C | 847 | 40 | 16.91 | 17.38 | 8.65 |
| Zyxin | 728 | 35 | 20.26 | 21.59 | 12.90 |
| VASP | 530 | 35 | 36.83 | 37.89 | 13.21 |

3) Gelatin
| Protein | N<sub>FA</sub> | N<sub>cells</sub> | MedianZ<sub>FA</sub>(nm) | MeanZ<sub>FA</sub>(nm) | Std.dev. (nm) |
| --- | --- | --- | --- | --- | --- |
| ILK | 919 | 50 | 0 | 1.74 | 9.89 |
| FAK | 801 | 50 | 5.89 | 6.48 | 11.18 |
| Paxillin | 757 | 50 | 5.16 | 5.68 | 11.42 |
| Talin-N | 155 | 50 | 2.61 | 2.52 | 5.71 |
| Talin-C | 781 | 50 | 12.91 | 15.87 | 15.72 |
| Vinculin-N | 273 | 50 | 15.93 | 16.50 | 8.21 |
| Vinculin-C | 535 | 50 | 35.29 | 35.43 | 10.35 |
| Zyxin | 439 | 50 | 62.75 | 61.28 | 17.09 |
| VASP | 379 | 50 | 63.57 | 64.77 | 15.98 |

As shown in Fig. 7A-B, we found that mESC FAs on gelatin and laminin exhibit largely similar hierarchical organization as on fibronectin, with ILK, paxillin, and FAK in the ISL, talin in an oriented organization defining a similarly compact (∼10 nm) FTL, and zyxin and VASP in the ARL. Interestingly, while on gelatin vinculin was located largely within the ARL (Fig. 7A, Table 1), we noticed that on laminin vinculin exhibited a significant FTL localization. Furthermore, unlike on fibronectin or gelatin or in “classical” FAs, where vinculin was oriented with the C-terminus upward, on laminin the vinculin C-terminus was lower than its N-terminus, indicative of an opposite molecular orientation. In conjunction, the ARL zone on laminin appeared to be relatively restricted in range. Taken together, these results suggest that while the multilayer stratification of FAs appears to persist across different ECMs, different integrin-ECM signaling could participate in the modulation of the nanoscale distribution of a subset of FA proteins, particularly vinculin. The interactions that underlie these observed changes are currently unknown, since the C-terminal domain of vinculin contain binding sites to multiple partners including F-actin, paxillin, PIP2, and can be dimerized or phosphorylated by Src^57^. How these interactions are regulated in response to different ECMs are still not fully understood and awaits further investigation.

### Discussion

In this study we characterized the integrin adhesions-mediated process in mESCs at the molecular and nanostructural level, motivated by the attenuated mechano-responses of mESCs^20-21^. Our findings reaffirmed that adherent mESCs exhibit subdued responses to substrate rigidity cues in significant contrast to typical fibroblastic or mesenchymal responses, consistent with previous reports^21^. The average spread area of mESCs was ∼450 μm^2^ and remains largely similar over the range of 0.6 - 55 kPa (Fig. 3). In conjunction, the increasing stiffness of the substrate appear to elicit no significant increase in stress fibers formation in mESCs, while FAs remained sparsely distributed and confined to the cell periphery. Nevertheless, despite the paucity and diminutive morphology of mESC FAs (Fig. 2), our analysis reveals that the majority of the core components of the FAs are expressed at the transcriptome level, localized to FAs at the protein level, and organized into the conserved multi-layer architecture at the nanostructural level. Furthermore, mESC FAs contain markers of mature FAs such as zyxin, and disassemble following inhibition of contractility by blebbistatin or Y-27632, indicative of their dependence on myosin II contractility.

Our results also suggest a number of interesting directions for further in-depth investigation at the molecular and nanostructural levels. First, the comparatively lowered expression of LIM domain proteins in PSCs corroborates previous proteomics analysis that documented their contractility-dependent enrichment in “classical” FAs^42^. The underlying molecular mechanisms of how LIM domain proteins participate in mechanotransduction are still not fully understood but may be potentially important since LIM domain proteins are known to shuttle into the nucleus to participate in transcription regulation^58-59^. Secondly, the variable nanoscale distribution and molecular orientation observed for vinculin further highlight its role as the “adjustable” mechano-elements within the otherwise conserved multi-layer FA architecture^27^. Our results implicate integrin-ECM signaling as a potential key input regulating vinculin configuration. Interestingly, our recent study in cadherin-based adhesions have documented the variable nanoscale organization, conformation, and orientation of vinculin as well^35^. However, vinculin in cadherin-based adhesions interacted with α_E_-catenin instead of talin and any commonality in their regulatory mechanisms has not been explored in details. In short, it remains an open question whether the relationship between the observed nanoscale organization of vinculin are ultimately related to mESC differentiation. We note that in hMSCs, recent studies have demonstrated the involvement differentiation of mechanosensitive binding of vinculin to MAPK1^60^, and thus analogous mechanisms could be in play in mESCs. Thirdly, while the attenuated mechanobiological responses of mESCs are distinctive of its naïve pluripotent states, once differentiation is committed these cells have been shown to ramp up their mechano-responses rapidly^61^. The concerted changes in transcription and cellular organization during this process could serve as an experimental platform to dissect fundamental mechanobiological processes.

We note that a recent pre-print has also reported a study of FA nanoscale architecture in human induced PSCs (hiPSCs)^62^. In that study, multi-layer stratification of FA proteins into ISL/FTL/ARL was observed, supporting our notion that the multi-layer architecture may be a conserved architecture of cell adhesion complexes^28^. Interestingly, on vitronectin ECM used in that study, a relatively high position of vinculin was also observed similar to our observation for mESC FAs on fibronectin and gelatin, while the inverted position of vinculin is similar to our mESC results on laminin. Altogether, this provides complementary findings to our results that vinculin appears to be the variable molecular components of the FA molecular clutch^15, 35^, although the molecular mechanisms involved remain unclear. We also note that the size of FAs in hiPSC are remarkably large (and larger than the parental fibroblasts) in that study. We note that these FAs appear to be unique to the edge cells of multi-cellular colonies, however, while interior cells have much less prominent FAs or actin cytoskeletal organization as observed previously^43^. These differences in FA profiles may arise from a combination of the differences between single cell and multi-cellular colony, cells in the edge and interior colony^63^, and the difference between the mESCs which is generally considered to correspond to ground-state pluripotency in comparison to the primed pluripotent states of hESCs for which the hiPSCs presumably recapitulate^64-65^.

Collectively, our data suggest that the basic molecular machinery of FAs appear to be operational in mESCs. We propose that such mESC FAs could perhaps be considered as “baseline” cell adhesion complexes, in contrast to the “classical” FAs that arise in later developmental stages which may have gained robust maturation and probably additional signaling capabilities (e.g. LIM domain protein-mediated pathways) during differentiation. In other words, while mESCs could sense and adhere to ECM, mechanoresponsive machineries which require the formation of pro-migratory and pro-contractility structures such as lamellipodia and dorsal stress fibers are yet to be mobilized. As these “positive feedback” aspects of the mechanosensitive responses appear to be rapidly upregulated following the exit from naïve pluripotency, further system-level analysis of transcription during differentiation could potentially yield deep insights into how the adhesion and cytoskeletal morphological profiles of the cells are regulated.

## Conclusion

Our study characterized how mESC FAs are spatially organized to perform their functions. Despite small sizes and peripheral distribution, mESC FAs exhibited a compositional signature of mature FAs and respond to myosin II contractility. At the nanoscale level, mESC FAs possess the core integrin-talin-actin connections and retain the multi-layer organization but with ECM-dependent variation in vinculin configuration. Our findings also highlighted interesting directions for further investigation of mESC mechanotransduction: i) the involvement of LIM domain proteins and ii) the ECM-dependent roles of the nanoscale configuration of vinculin.

## Acknowledgement

We thank Cheng Gee Koh (Nanyang Technological University, Singapore) for generous sharing of reagents and insightful discussion, and Sergey Plotnikov (University of Toronto, Canada) for technical assistance. We acknowledge funding support from the Singapore National Research Foundation (NRF) under the NRF Fellowship scheme (NRF-NRFF-2011-04), the NRF Competitive Research Programme (NRF2012NRF-CRP-001-084), the Ministry of Education Academic Research Fund Tier 2 (MOE-T2-1-124 and MOE-T2-1-045), and MBI seed funding to P.K. and E.Y. S.X. was also supported in part by the MBI Graduate Scholarship.

---

**Figure S1.**
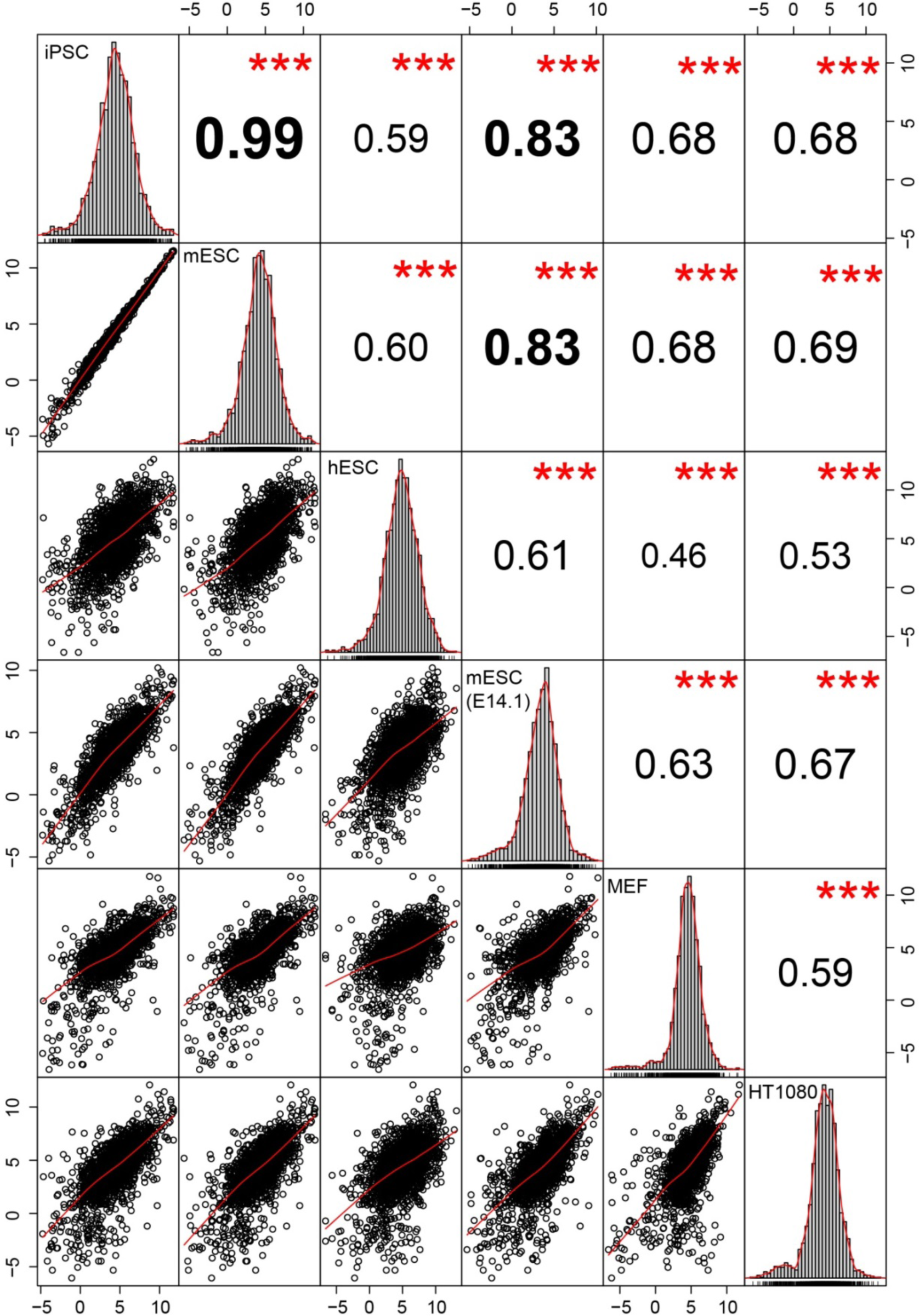
Meta-adhesome correlation Chart. Gene expression profiles (Log_2_ FPKM) of 1999 meta adhesome for 6 cell lines were plotted in a Pearson correlation chart. The diagonal shows the gene expression distribution of each cell lines. The bivariate scatter plots with fitted line were displayed below the diagonal, and the value of correlation coefficients with the significance level as stars for Pearson correlation were presented above the diagonal. ***: P<0.001.

**Figure S2.**
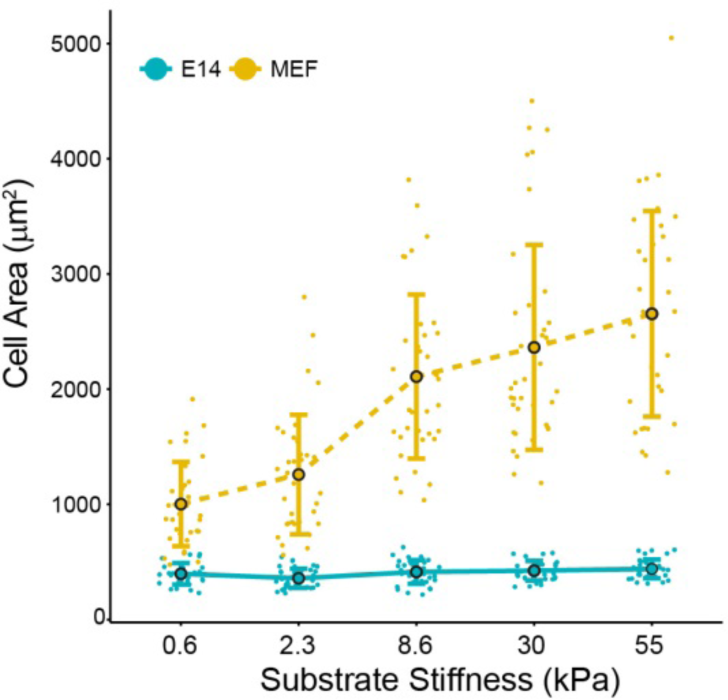
Cell spreading area on substrate with various rigidity. Quantifications of cell spreading area as shown in **Figure 2 (C)**. Data are presented as mean with SD. Each data point represents a cell (n_cell-E14_=31, 34, 34, 32, 36 and n_cell-MEF_=33, 37, 37, 39, 34 for 0.6, 2.3, 8.6, 30, 55 kPa).

**Figure S3:**
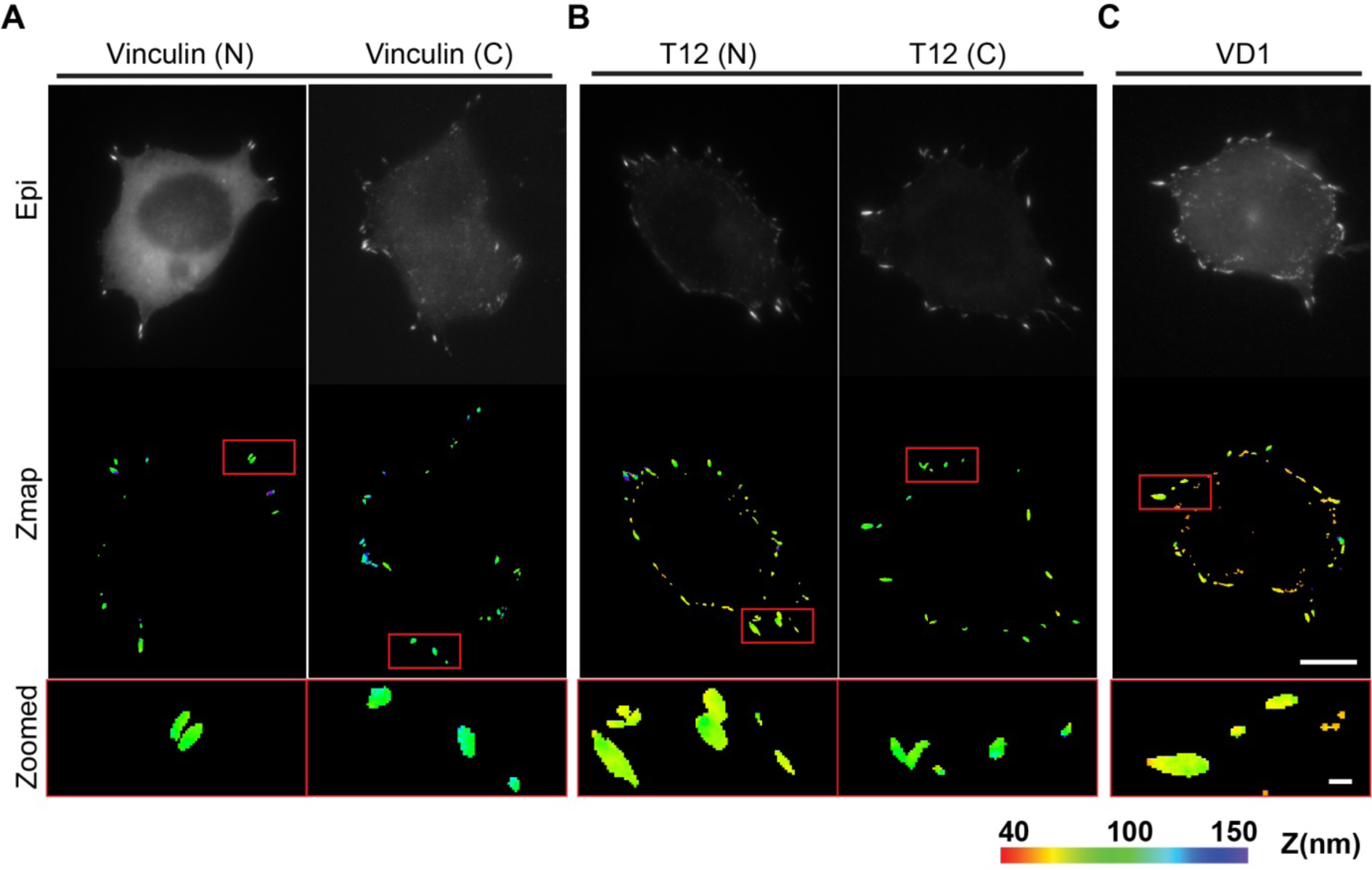
Topographic map of the z-position for vinculin variants expressed in mESCs on fibronectin. **(A-C)** Representative images of Epi and color-encoded map for Vinculin-N, Vinculin-C **(A)**, T12-N, T12-C **(B)** and VD1**(C)**, related to **Figure 6 (D)**. Boxed regions were magnified in below. Scale bars: 10 μm; zoomed: 1 μm. Color bar indicates Z-position relative to the surface.

**Figure S4.**
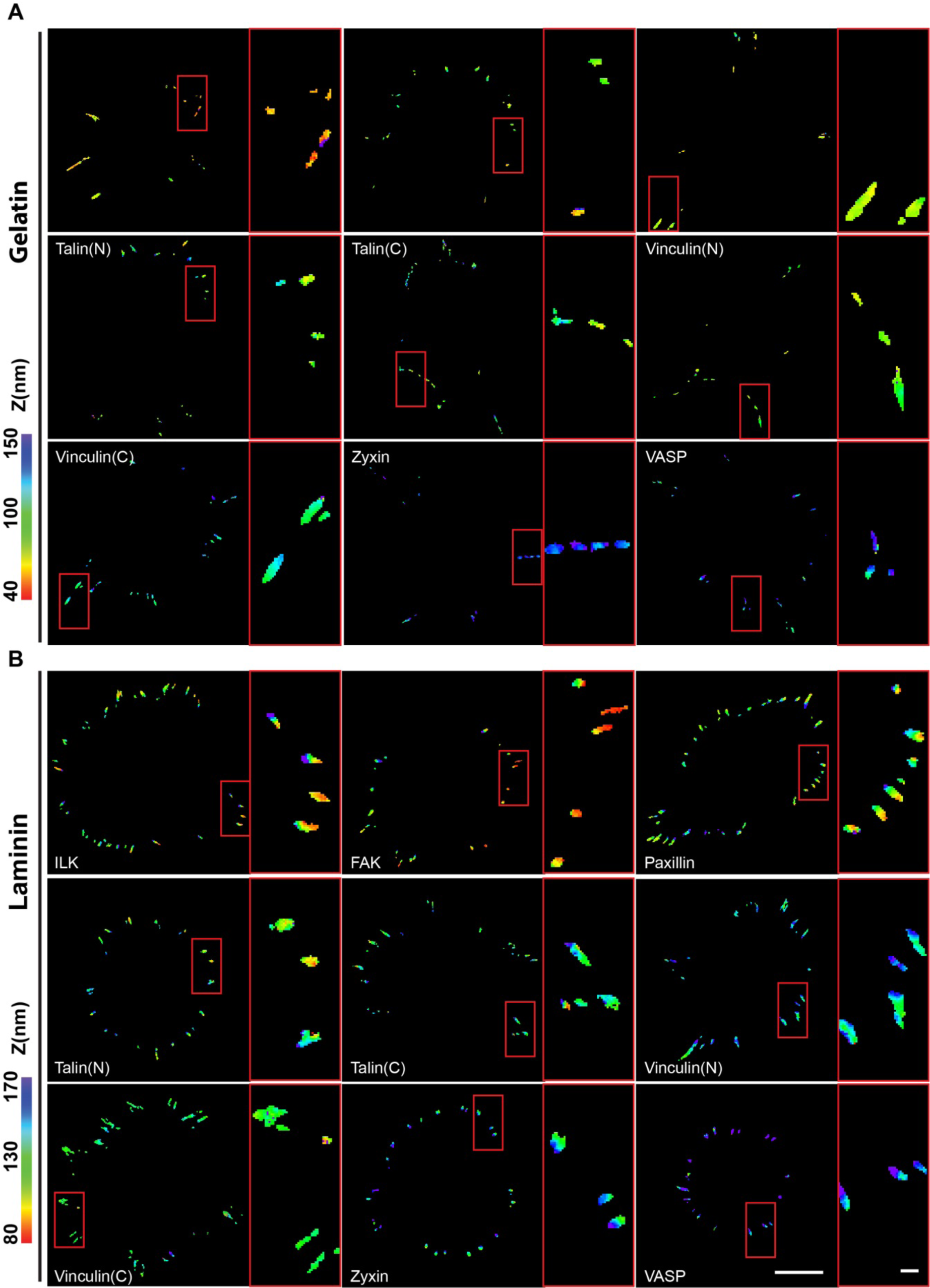
Topographic map of the z-position for integrin adhesome proteins in mESCs on gelatin and laminin. **(A, B)** Representative topographic map of FA proteins on gelatin **(A)** and laminin **(B)** with color-encoded Z-position as measured by surface-generated structured illumination microscopy, related to **Figure 7**. The magnified view of each boxed region was shown at the right side of each panel. Color bar indicates the Z relative to the surface. Scale bar: 10 μm; inset: 1 μm.

